# System-Specific Epigenetic Aging Signatures in Autistic Adults

**DOI:** 10.64898/2026.09.12.751174

**Authors:** Abigail Dickinson, Candace R. Lewis, Daniel H. Geschwind, Judith E. Carroll, Catherine Lord

**Affiliations:** Department of Psychiatry and Biobehavioral Sciences, Jane and Terry Semel Institute for Neuroscience and Human Behavior, David Geffen School of Medicine at UCLA, University of California, Los Angeles; Center for Autism Research and Treatment, Jane and Terry Semel Institute for Neuroscience and Human Behavior, University of California, Los Angeles; School of Life Sciences, Arizona State University; Department of Psychology, Arizona State University; Cousins Center for Psychoneuroimmunology, Jane and Terry Semel Institute for Neuroscience and Human Behavior, University of California, Los Angeles; Department of Education, School of Education and Information Studies, University of California, Los Angeles; Department of Neurology, David Geffen School of Medicine at UCLA, University of California, Los Angeles; Department of Human Genetics, David Geffen School of Medicine at UCLA, University of California, Los Angeles

**Author notes:** Correspondence concerning this article should be addressed to Abigail Dickinson, Department of Psychiatry and Biobehavioral Sciences, Semel Institute for Neuroscience and Human Behavior, University of California, Los Angeles, Los Angeles, CA, USA.

**Keywords:** autism, epigenetic aging, DNA methylation, SystemsAge, quality of life, psychosocial factors

## Abstract

**Background:** Autism is a lifelong neurodevelopmental condition, but the biological processes shaping aging are poorly understood. DNA methylation-based measures index physiological decline and mortality risk independently of chronological age. We tested whether biological aging differs in autistic adults, whether differences are system-specific, and which factors predict within-autism variation.

**Methods:** Saliva DNA methylation (Illumina EPIC) was assayed in 37 autistic adults (31 male, 6 female; mean age 29.8 years), followed from early childhood, spanning a broad ability range including intellectual disability, and 188 non-autistic adults from four public datasets, processed jointly. Fifteen measures were calculated: four global (Horvath, GrimAge, DunedinPACE, SystemsAge) and 11 system-specific SystemsAge subscores. Models adjusted for age, sex, and epithelial-cell proportion.

**Results:** Global aging measures did not differ between groups (all q > .34). However, autistic adults had higher Brain (β = 0.81), Blood (β = 0.65), and Liver SystemsAge (β = 0.58; all q < .01), with Brain and Blood robust across sensitivity analyses. Within autism, aging measures were unrelated to symptom severity, IQ, co-occurring diagnoses, or medication use. Instead, elevations were associated with psychosocial circumstances, including a lack of structured daytime activity, greater loneliness, and lower quality of life.

**Conclusions:** Differences in aging biology in autism appeared system-specific rather than global, with no evidence of generalized epigenetic age acceleration. This vulnerability appeared more closely tied to psychosocial factors than core clinical features. System-specific DNAm measures may therefore provide a more sensitive framework for identifying biological heterogeneity and potentially modifiable contributors to healthy aging in autistic adulthood.

## 1. Introduction

Autism is a lifelong neurodevelopmental condition, yet research has focused disproportionately on childhood and early development. As a result, relatively little is known about how autistic people age, including how biological aging unfolds, and which environmental and psychosocial factors shape variability in health and functioning across adulthood (1). Although research interest in middle-aged and older autistic adults has increased, this work remains limited and continues to underrepresent individuals with intellectual disability or greater support needs. Addressing this gap is increasingly urgent as improved recognition across the lifespan and broader demographic aging contribute to a growing population of older autistic adults, and as emerging evidence indicates that rates of age-related disease are elevated in this group (2,3). Understanding biological aging in autism is therefore critical for identifying vulnerability to age-related disorders, guiding efforts to support autistic adults as they age, and elucidating potentially modifiable risk factors that shape aging trajectories.

Existing research using health, cognitive, and neural markers provides mixed evidence for altered aging trajectories in ASD. Population-level studies report elevated rates of cardiovascular, metabolic, and neurodegenerative conditions (2–4), alongside some evidence of reduced life expectancy (5,6). In contrast, findings from cognitive and neural markers are less consistent. Although accelerated aging has been reported in some domains (7,8), most findings suggest broadly similar age-related patterns in autistic and non-autistic individuals, with little consistent evidence of disproportionate decline in memory, processing speed, executive function, or structural and functional brain measures (for reviews, see (9–12)).

Molecular biomarkers offer a complementary approach by indexing biological differences that may precede overt clinical change. DNA methylation (DNAm)-based biomarkers, including epigenetic clocks, are among the most widely studied molecular measures of biological aging. These algorithms use age-associated methylation patterns across the genome to index different aspects of aging and are associated with morbidity, functional decline, and all-cause mortality independently of chronological age (13,14). In autism, such measures may help identify biological vulnerability and determine whether it is concentrated in specific biological processes or physiological systems. Because aging is not uniform across the body, successive generations of DNAm measures trained on increasingly specific outcomes provide complementary information at multiple levels.

First-generation clocks, such as the Horvath clock, primarily predict chronological age and capture broad age-related methylation patterns (15). Later-generation measures target more clinically relevant features: PhenoAge reflects phenotypic risk, GrimAge emphasizes mortality-related risk, and DunedinPACE estimates the pace of physiological decline (16–18). More recently, system-specific approaches have been developed. SystemsAge, for example, uses methylation-based surrogates of clinical and functional biomarkers to estimate mortality-related aging across eleven physiological systems (19). Applied together, these measures make it possible to examine whether aging-related differences are global or concentrated within biological systems.

Applying global and system-specific measures together may help clarify biological aging processes in autism. Prior work using first- and second-generation clocks has found limited evidence of global age acceleration (Gentilini et al., 2023; Liu et al., 2023; Okazaki et al., 2022), but whether more specific measures would reveal differences in particular physiological systems remains to be determined. A similar pattern has been observed in schizophrenia, where first-generation clocks yielded null findings, whereas second-generation and system-specific measures identified alterations in specific physiological domains (20–23).

Building on this work, we applied a broad panel of DNAm measures to characterize biological aging in autism at both global and system-specific levels. The panel included measures trained on chronological age, phenotypic and mortality risk, pace of aging, and aging within specific physiological systems. We applied these measures in a deeply-phenotyped cohort of autistic adults diagnosed early in life, followed for decades, and repeatedly evaluated with expert clinical assessment. The cohort spanned a broad range of cognitive ability and included individuals with intellectual disability, who are absent from prior epigenetic aging studies. Autistic and non-autistic methylation data were processed jointly, and the principal findings were tested across a series of sensitivity analyses. We first tested for group differences across the panel and then examined whether variation within the autistic group was associated with clinical, health, cognitive, and psychosocial characteristics, with the aim of identifying group-level and individual patterns of biological vulnerability to guide future longitudinal research on healthy aging in autism.

## 2. Methods

### 2.1. Participants

#### 2.1.1. Autistic Participants

Autistic participants were drawn from the Early Diagnosis Study (EDX), a longitudinal cohort recruited through consecutive referrals for autism evaluation before 3 years of age and followed into adulthood (24–26). The present analyses include 37 autistic adults (31 male, 6 female; mean age = 29.8 years, SD = 1.1, range = 27.6–32.8) who provided saliva during the age-30 assessment wave, which was disrupted by the COVID-19 pandemic and therefore captured a subset of the broader cohort.

All participants had a confirmed best-estimate clinical diagnosis of autism spectrum disorder (Section 2.2.1), and the sample spanned a broad range of cognitive and language abilities, including intellectual disability (Table 1). Procedures were approved by the UCLA Institutional Review Board. Participants with capacity provided written informed consent; assent and guardian consent were obtained where appropriate.

#### 2.1.2. Control Participants

Control DNAm data were drawn from four independent datasets in the NCBI Gene Expression Omnibus (GEO; GSE151485, GSE232332, GSE111631, GSE232891). Datasets were selected using prespecified criteria for tissue type, participant age, array platform, and metadata availability (see Methods S1). Raw IDAT files from all eligible control participants (n = 204) were processed together with the autistic samples using the same pipeline, retaining 188 after quality control. Sample characteristics are reported in Table 1 and, by dataset, in Table S1.

### 2.2. Clinical Assessments (Autistic Participants Only)

Unless otherwise specified, analyses used the in-person assessment closest to saliva collection, and the interval between assessment and saliva collection was included as a covariate where relevant. Measure-specific sample sizes and assessment ages are reported in Table 2.

#### 2.2.1. Autism Diagnosis

All 37 participants had a best-estimate clinical diagnosis established by expert clinicians using the ADOS-2 (27), the ADI-R (28), clinical judgment, following established procedures (29). Most were diagnosed in early childhood; diagnostic timing is reported in Table 2 and Methods S2.

#### 2.2.2. Physical Health, Co-occurring Conditions, and Medication Use

Body mass index (BMI; kg/m²) was calculated from measured height and weight, available for 34 of 37 participants, and categorized as <25 versus ≥25 kg/m² for descriptive analyses. Co-occurring conditions were characterized from the age-25 wave onward, available for 36 of 37 participants, and grouped into broad psychiatric, neurological, and physical health categories using prespecified coding rules (Methods S2).

Intellectual disability was examined separately through IQ-based subgroup analyses and was not treated as a co-occurring condition. Medication exposure was coded as any history of centrally acting psychotropic or neuroactive medication use during adulthood (Methods S3).

#### 2.2.3. Autism Symptomatology and Cognitive Ability

Autism symptom severity was indexed using ADOS-2 Calibrated Severity Scores for Social Affect and Restricted and Repetitive Behaviors (27). Cognitive ability was assessed using a developmentally appropriate hierarchical battery (30), and participants were grouped by verbal IQ (VIQ ≥ 70 vs. VIQ < 70) for descriptive and exploratory analyses.

#### 2.2.4. Psychosocial Functioning and Wellbeing

Structured daytime engagement was coded according to whether participants regularly participated in paid or voluntary employment, formal education, vocational training, day programs, or supported daytime programming. Loneliness was assessed using the 24-item Asher Loneliness and Social Dissatisfaction Scale (31). Quality of life was assessed using two global items from an adapted WHOQOL-BREF (32,33), rated 1 to 5, indexing overall quality of life and satisfaction with overall health; self-and caregiver-report data were available for 28 participants. Procedures for selecting among multiple available ratings are described in Methods S2.

### 2.3. DNA Methylation Data

#### 2.3.1. Data Acquisition

Saliva was collected using Oragene DNA self-collection kits (DNA Genotek, Ottawa, ON, Canada). Genomic DNA was extracted according to manufacturer protocols, bisulfite converted, and assayed on the Illumina HumanMethylationEPIC v1.0 BeadChip (Illumina, San Diego, CA, USA). Control datasets were saliva-derived and generated on the same platform (Table S1).

#### 2.3.2. Preprocessing and Quality Control

Autistic and control data were processed jointly in R using established pipelines for saliva-derived methylation data (34–37). Samples were excluded for excessive probe-detection failure, low signal intensity, low bead count, or sex discordance; 16 samples, all from control datasets, were excluded, yielding 37 autistic and 188 control participants. CpGs were excluded for poor detection, low bead count, sex-chromosome location, or SNP overlap, leaving 810,952 for primary analyses, with cross-reactive probes additionally excluded in sensitivity analyses (38). Data were normalized using functional normalization (39), and cell-type composition was estimated using EpiDISH with an EPIC-based saliva reference panel (40), with estimated epithelial-cell proportion included as a covariate. Full thresholds, probe-removal counts, and software implementation are provided in Methods S4 and Table S2.

### 2.4. Epigenetic Aging Measures

Analyses focused on 15 DNAm aging measures spanning global and system-specific aspects of biological aging, calculated from quality-controlled β-values. Global measures included the Horvath skin-and-blood clock (41), GrimAge (18), DunedinPACE (16), and the SystemsAge composite, which additionally yields 11 domain-specific subscores capturing aging-related variation across physiological systems (19); these are referred to by their system labels (e.g., Brain SystemsAge). Methylation-derived scores for BMI (42), smoking (43), and general cognitive ability (44) were calculated for use as covariates. Implementation details and probe coverage are provided in Table S3.

### 2.5. Statistical Analyses

Analyses were conducted in R 4.5.1. Between-group demographic differences were assessed using t-tests and chi-square or Fisher’s exact tests as appropriate (Table 1). Unless otherwise specified, epigenetic outcomes were analyzed using linear regression adjusted for chronological age, sex, and estimated epithelial-cell proportion, with outcome-specific extreme observations excluded when residuals fell more than 3 interquartile ranges beyond the first or third quartile of a covariate-only model. Regression assumptions were evaluated using residual and influence diagnostics. Benjamini–Hochberg FDR correction was applied within prespecified analysis families. All tests were two-sided; p < .05 was considered nominally significant and q <.05 significant after correction.

#### 2.5.1. Associations with Chronological Age

Associations with chronological age were examined for the three measures intended to index the level of biological aging (Horvath, GrimAge, SystemsAge composite) in the combined sample and descriptively within the autistic subgroup. SystemsAge subscores and DunedinPACE were not included because they do not estimate chronological age.

#### 2.5.2. Group Differences in Epigenetic Measures

Group differences were tested for each of the 15 measures, standardized across the full analytic sample so that coefficients represent adjusted differences in standard-deviation units, with positive values indicating higher scores in the autistic group. To assess robustness to preprocessing and residual technical variation, analyses were repeated using noob background-corrected data generated in minfi, with the first 10 technical principal components derived from control-probe intensities using meffil.

#### 2.5.3. Sensitivity Analyses

Sensitivity analyses evaluated robustness to sample composition, probe filtering, preprocessing, additional DNAm-derived covariates, and the influence of individual control datasets, comprising an age- and sex-restricted subsample, exclusion of cross-reactive probes, adjustment for DNAm BMI, smoking, and cognitive ability, and leave-one-control-dataset-out analyses.

#### 2.5.4. Within-Autism Analyses

Within-autism analyses focused on the four SystemsAge subscores showing group-level elevations (Brain, Blood, Liver, Metabolic). Associations with sociodemographic, clinical, health, cognitive, and psychosocial characteristics were examined using linear regression adjusted for chronological age, sex, and estimated epithelial-cell proportion, with DNAm BMI added in secondary models and the assessment-to-saliva interval added where predictor timing differed. Categorical comparisons were restricted to groups with at least five participants and, where necessary, collapsed into binary categories (Methods S5). Given the modest sample size and exploratory nature of these analyses, findings were interpreted with emphasis on effect sizes, confidence intervals, and consistency across models.

## 3. Results

### 3.1. Age Associations

All three epigenetic measures intended to estimate biological age were positively associated with chronological age in the combined sample (all p < .001; Figure 1), most strongly for the Horvath skin-and-blood clock (r = .94), followed by GrimAge (r = .87) and the SystemsAge composite (r = .65). Associations were also positive within the autistic subgroup, although less precise given the smaller sample and restricted age range (Results S1).

**Figure 1.**
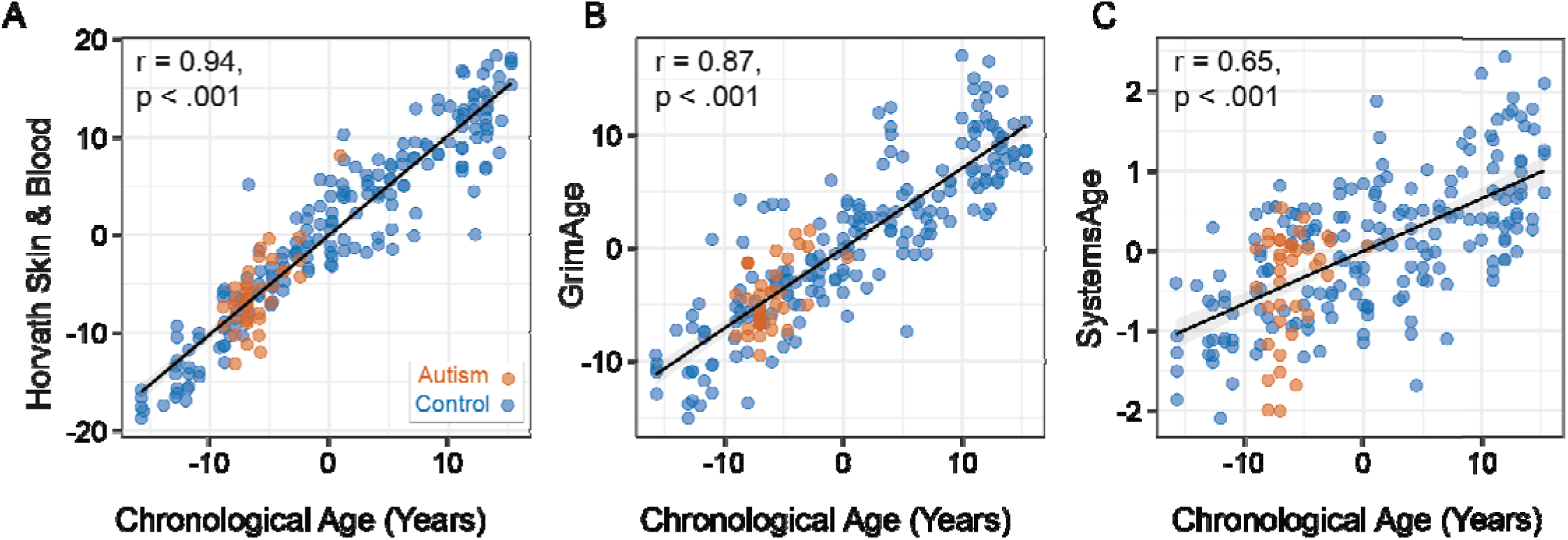
Associations between chronological age and DNA methylation–derived aging measures. Partial associations between chronological age and (A) Horvath skin-and-blood age, (B) GrimAge, and (C) the SystemsAge composite, adjusted for sex and estimated epithelial-cell proportion. Axes show covariate-adjusted residuals; solid lines show fitted associations with 95% confidence bands. Points represent individual participants and are colored by diagnostic group (autistic = red; control = blue).

### 3.2. Group Differences

Primary analyses compared epigenetic aging measures between autistic adults (n = 37) and non-autistic controls (n = 188), adjusting for chronological age, sex, and estimated epithelial-cell proportion (Figure 2; Table 3). Autistic adults had higher scores on three system-specific SystemsAge measures, all surviving FDR correction across the full 15-measure panel: Brain (β = 0.81, q < .001), Blood (β = 0.65, q = .002), and Liver SystemsAge (β = 0.58, q = .008). Metabolic SystemsAge was nominally higher but did not survive correction (β = 0.38, p = .035, q = .130), and the remaining seven subscores did not differ (all q > .26). None of the four composite measures differed between groups: Horvath, GrimAge, DunedinPACE, and the SystemsAge composite (all q > .34).

**Figure 2.**
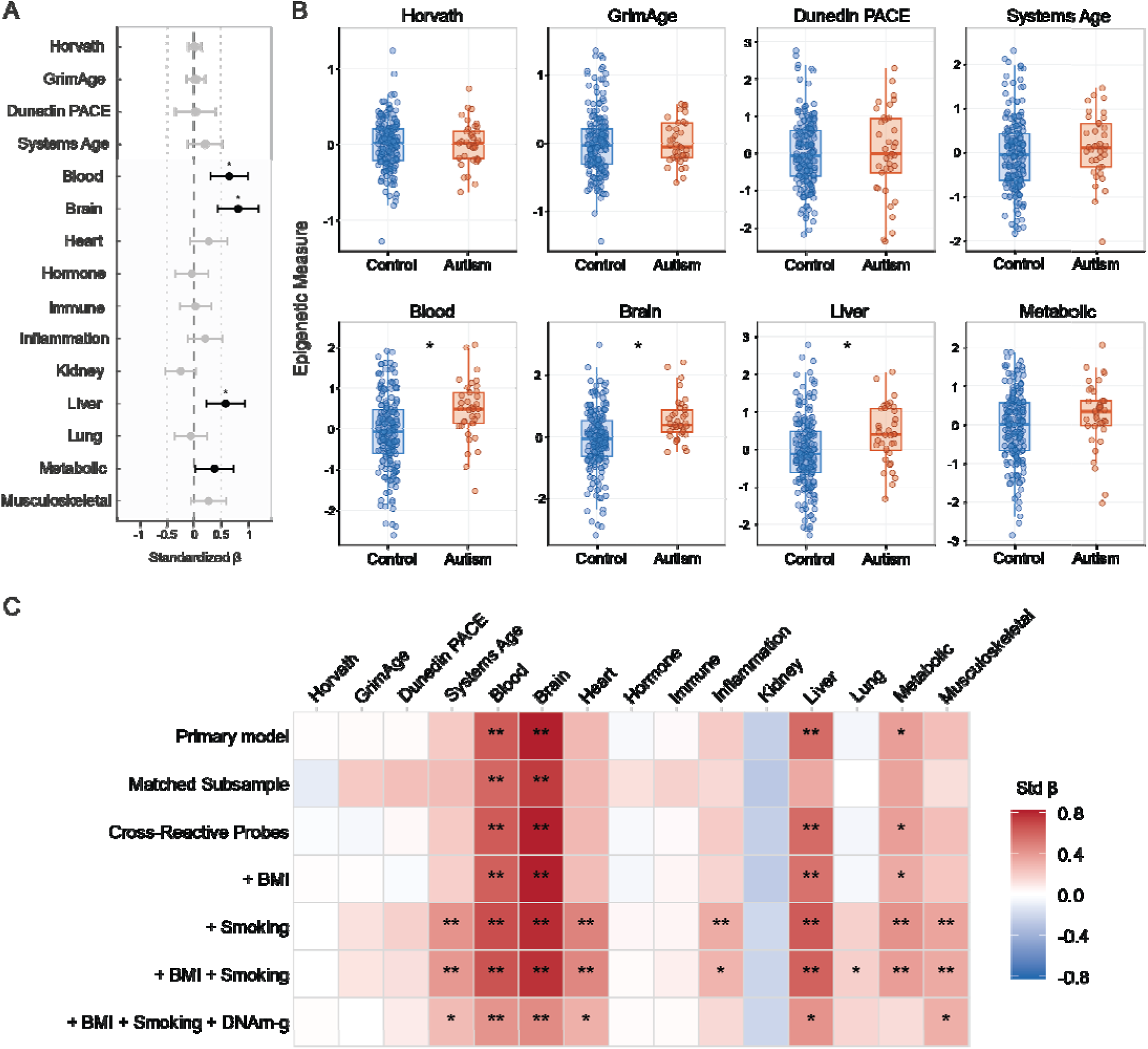
Group differences in DNA methylation–derived aging measures. (A) Forest plot of adjusted autism–control differences across all 15 measures, expressed in standard-deviation units. Positive coefficients indicate higher scores in autistic participants. Black markers indicate nominal significance (p < .05), and asterisks indicate FDR significance (q < .05). (B) Distributions of covariate-adjusted scores for the four global/composite aging measures (upper row) and the four SystemsAge subscores showing nominal or FDR-significant elevations in autistic adults (lower row). (C) Heatmap of standardized group-difference coefficients across the primary model and six sensitivity models evaluating sample composition, cross-reactive probe exclusion, and additional adjustment for DNAm BMI, DNAm smoking, and DNAm-g. Red indicates higher scores in autistic participants and blue indicates lower scores. **q < .05; *p < .05.

### 3.3. Sensitivity Analyses

Brain and Blood SystemsAge differences were highly consistent across sensitivity analyses (Figure 2C; Table S4), remaining FDR-significant in the age- and sex-restricted sample, after exclusion of cross-reactive probes, and after adjustment for DNAm BMI and smoking. Liver SystemsAge was elevated across most specifications, although evidence was weaker in some models, whereas the Metabolic difference reached significance only under selected covariate specifications. Composite measures remained nonsignificant throughout. Leave-one-dataset-out analyses indicated that the Brain, Blood, and Liver differences were not attributable to any single control dataset, remaining positive and FDR-significant after exclusion of each dataset in turn (Figure S1). The Metabolic difference was less stable across iterations, indicating greater sensitivity to control-sample composition.

### 3.4. Within Autism Correlates of SystemsAge

To characterize individual variation in epigenetic aging within the autistic sample, we examined associations between the four SystemsAge subscores showing group-level elevations and a range of demographic, clinical, and psychosocial factors. We first evaluated the relationship between BMI calculated from measured height and weight (hereafter, BMI) and a methylation-derived measure of BMI (DNAm BMI). The two were strongly correlated (r = .72, p < .001; Figure 3A). Higher BMI was associated with higher Brain (partial r = .41, p = .017) and Metabolic SystemsAge (partial r = .42, p = .013), and DNAm BMI was associated with all four subscores (Figure 3B–C; Table S5). Within-autism analyses were therefore repeated with DNAm BMI as an additional covariate.

**Figure 3.**
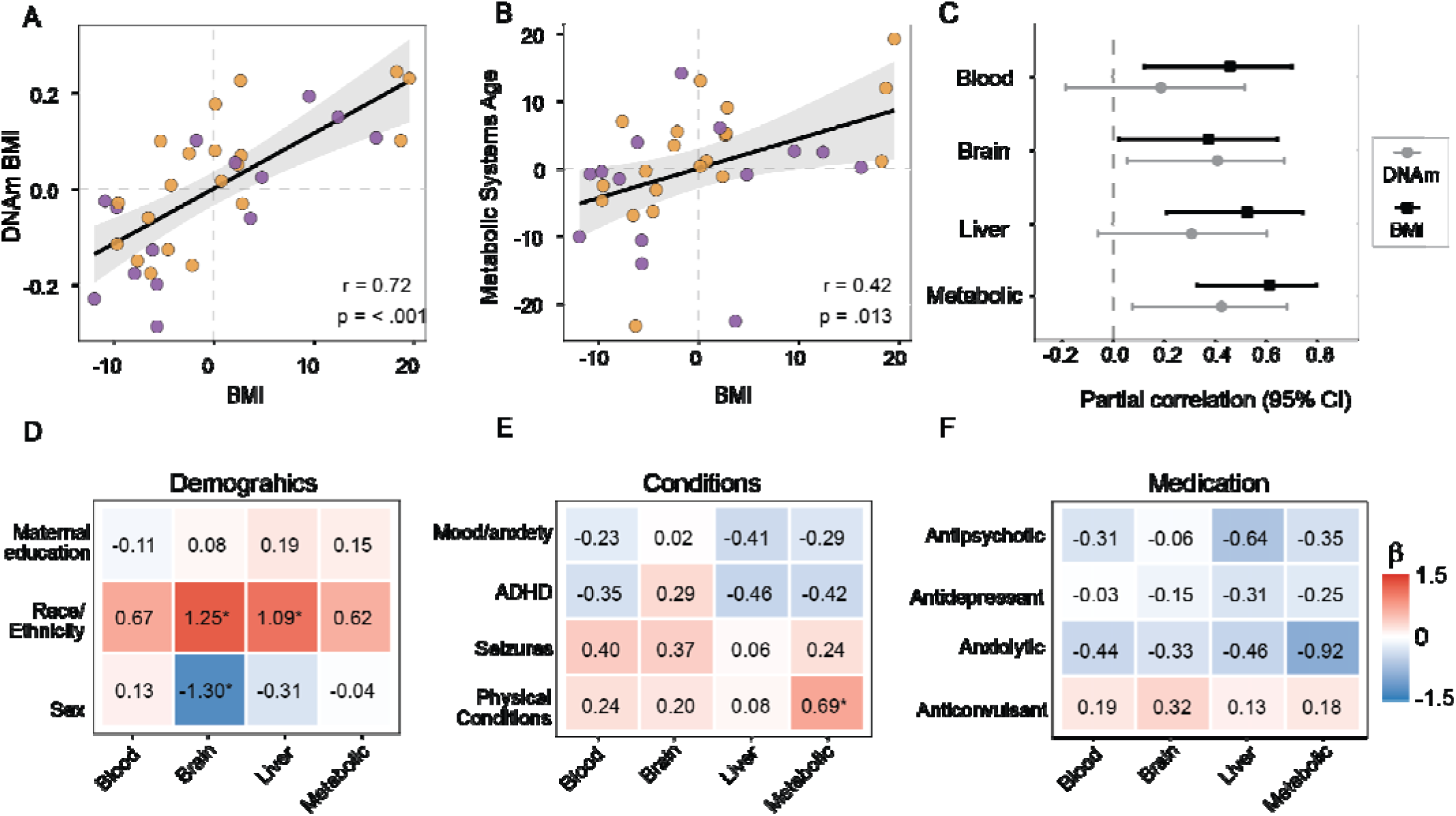
Associations between BMI and SystemsAge-related characteristics within the autistic sample. (A) Association between measured BMI and DNAm BMI. (B) Association between measured BMI and Metabolic SystemsAge. (C) Forest plot comparing partial correlations of measured BMI and DNAm BMI with Blood, Brain, Liver, and Metabolic SystemsAge. Points represent partial correlation coefficients and horizontal lines represent 95% confidence intervals. (D–F) Heatmaps of standardized effect sizes for associations between the four SystemsAge subscores and (D) demographic characteristics, (E) co-occurring conditions, and (F) medication exposure. In panel D, negative values indicate higher SystemsAge scores among participants who were non-Hispanic White, female, or whose mothers had more than 4 years of postsecondary education. In panels E and F, positive values indicate higher SystemsAge scores in the presence of the corresponding co-occurring condition or medication exposure. *p < .05.

#### 3.4.1. Demographic Factors

Racially and ethnically minoritized participants had higher Brain and Liver SystemsAge than non-Hispanic White participants, and females had higher Brain SystemsAge than males. All three associations persisted after adjustment for DNAm BMI. No other demographic comparisons were significant (Figure 3D; Table S6).

#### 3.4.2. Co-occurring Conditions and Medication Use

Co-occurring conditions and medication exposure showed few associations with SystemsAge (Figure 3E–F; Tables S7–S8). Participants with a co-occurring physical health condition had higher Metabolic SystemsAge in the primary model (β = 0.69, p = .020), but this was attenuated after adjustment for DNAm BMI (β = 0.36, p = .210). No other comparisons were significant.

#### 3.4.3. Autism Symptomatology and Cognitive Ability

Neither ADOS-2 calibrated severity scores nor full-scale, verbal, or nonverbal IQ were associated with the four SystemsAge subscores, and scores did not differ between participants with VIQ ≥ 70 and VIQ < 70 (Tables S6, S9).

**Figure 4.**
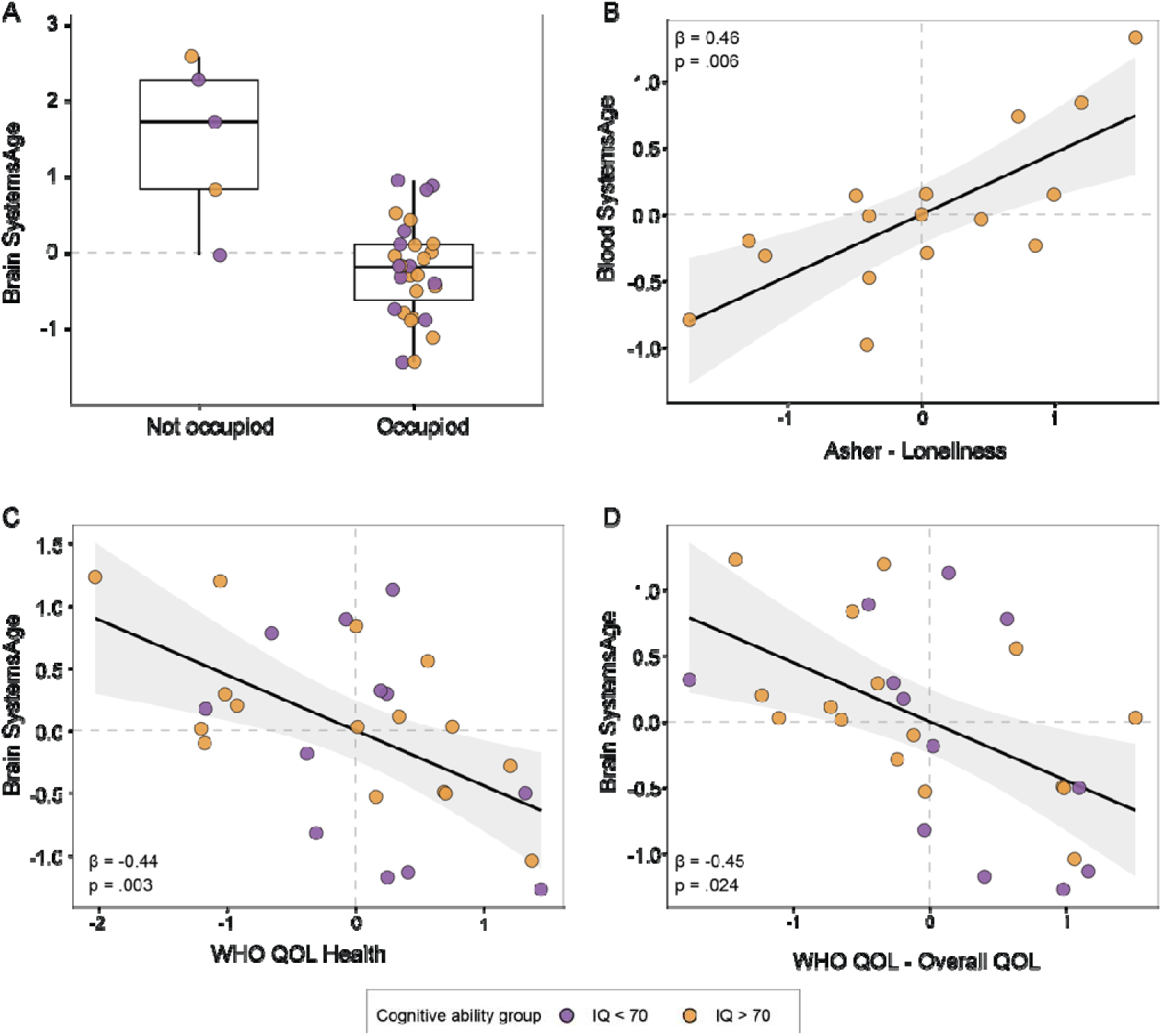
Associations between SystemsAge subscores and psychosocial characteristics within the autistic sample. (A) Differences in Brain SystemsAge according to structured daytime engagement, comparing participants who were currently engaged in structured daytime activity with those who were not. (B) Association between Blood SystemsAge and loneliness, assessed using the Asher Loneliness and Social Dissatisfaction Scale. (C–D) Associations between Brain SystemsAge and WHOQOL-BREF ratings of (C) overall health and (D) overall quality of life. Plotted values are residualized for chronological age, sex, estimated epithelial-cell proportion, and the interval between psychosocial assessment and saliva collection where applicable. Individual participants are shown as points and color-coded by verbal IQ group (VIQ ≥ 70, orange; VIQ < 70, purple).

#### 3.4.4. Psychosocial Characteristics

Participants not currently engaged in structured daytime activity showed higher SystemsAge than occupied participants. In models adjusted for age, sex, and estimated epithelial-cell proportion, the not-occupied group had higher Blood (β = 0.85, p = .011), Brain (β = 1.76, p = .003), and Metabolic SystemsAge (β = 1.15, p < .001), whereas the Liver difference did not reach significance (β = 0.96, p = .064). After additional adjustment for DNAm BMI and race and ethnicity, the Brain (β = 1.68, p = .005) and Metabolic associations (β = 0.83, p = .006) remained significant, while the Blood association was attenuated (β = 0.54, p = .092).

Lower WHOQOL-BREF ratings of overall quality of life and overall health were associated with higher Brain SystemsAge (β = −0.44, p = .024 and β = −0.43, p = .003), and both remained significant in fully adjusted models (Table S10). Lower overall quality of life was also associated with higher Metabolic SystemsAge (β = −0.48, p = .019), although this did not persist after additional adjustment. Caregiver-report sensitivity analyses yielded a similar pattern for Brain SystemsAge (Table S10).

Greater loneliness was associated with higher Blood SystemsAge (β = 0.54, p = .006), and this association remained significant after adjustment for DNAm BMI and for race and ethnicity (β = 0.59, p = .030).

## 4. Discussion

In this deeply phenotyped cohort of autistic adults followed from early childhood, we identified a selective pattern of system-specific epigenetic aging differences that was not apparent on conventional global measures. The clearest and most robust elevations were observed in Brain and Blood SystemsAge, and these findings persisted across multiple sensitivity analyses. In contrast, autistic adults did not differ from non-autistic controls on the Horvath skin-and-blood clock, GrimAge, DunedinPACE, or the overall SystemsAge composite. Within the autistic group, variation in SystemsAge was not strongly related to autism symptom severity or cognitive ability, but showed potentially meaningful associations with structured daytime engagement, quality of life, and loneliness.

These findings fit with a broader view of aging as a multidimensional process that does not occur uniformly across physiological systems, and they follow a pattern documented in other conditions. In schizophrenia, early studies using first-generation clocks reported largely null results (22,23), whereas later work using second-generation and system-specific measures revealed alterations in particular physiological domains (20,21), although the pattern there appears more widespread across SystemsAge domains than what we observed. A comparable divergence is emerging in autism. Prior studies using first- and second-generation clocks found limited evidence of global age acceleration (45–47), while more specific signals have appeared in particular tissues or subgroups. Liu et al. (2023), for example, reported no overall difference in Horvath epigenetic age alongside evidence of accelerated cerebellar aging in a subset of older autistic adults.

Differences in samples, tissues, and study design may contribute to these divergences, but the consistency of the pattern suggests that global measures can miss biologically meaningful variation. The selective profile observed here, concentrated in Brain and Blood SystemsAge, suggests that these elevations may reflect a distinct pattern of physiological vulnerability in autism rather than a nonspecific consequence of having a neurodevelopmental or psychiatric condition.

In contrast with measures trained on chronological age alone, such as the Horvath clock, Brain and Blood SystemsAge were trained on biological features with relevance to known health patterns in autistic adulthood. The Blood score incorporates hematological measures including indices of red blood cell and platelet function, whereas the Brain score incorporates biomarkers related to neurotrophic function, cerebrovascular health, and cognitive performance (Sehgal et al., 2025). Elevated scores therefore indicate methylation profiles associated with less favorable age-related neurological or hematological characteristics. This is broadly consistent with epidemiological evidence of elevated neurological, cerebrovascular, and related health conditions among autistic adults (2,4,48–51).

The systems aging scores associated with autism should not be interpreted as direct measures of organ aging or as representing inherited genetic risk. Methylation patterns are shaped by developmental biology, health status, environmental exposures, chronic stress, and accumulated physiological burden, and autistic adults may encounter these influences through multiple pathways, including co-occurring health conditions, social disadvantage, reduced access to care, and the demands of unsupportive environments. Epigenetic measures may therefore capture cumulative wear and tear arising from these interacting influences.

The substantial variation in SystemsAge within the autistic group provided an opportunity to examine factors that may contribute to this variation. Scores were not strongly associated with traditional clinical measures of the autism phenotype, including symptom severity, IQ, and intellectual disability grouping. They also did not appear to be driven by health-related factors and were not related to medication exposure or most co-occurring diagnoses.

BMI was a notable exception. Both measured BMI and DNAm BMI were related to several SystemsAge measures, consistent with the established importance of metabolic health for aging more generally (52). However, adiposity did not account for the principal group differences, as Brain and Blood elevations persisted after adjustment for DNAm BMI. Metabolic health therefore appears relevant to individual variation in aging-related biology within autism, but it does not provide a complete explanation for the system-specific differences observed between autistic and non-autistic adults.

Instead, our results pointed toward broader psychosocial factors as a potentially important source of variation. Participants without regular structured daytime engagement had higher Brain and Metabolic SystemsAge and, in the primary model, higher Blood SystemsAge than participants who were regularly occupied. Exploratory analyses suggested that this was not specific to employment or education: those engaged in work or education and those participating in other structured daytime activities showed broadly similar SystemsAge levels. This suggests that regular daytime engagement itself may be relevant, potentially through overlapping processes such as routine, social contact, cognitive stimulation, physical activity, and access to support. This may also help explain the absence of clear associations with IQ or symptom severity. Individuals with greater support needs may, in some contexts, have access to more structured environments that buffer against chronic stressors, whereas individuals who appear more independent may experience greater cumulative exposure to the demands of navigating environments poorly adapted to autistic needs. Although speculative, this possibility highlights the importance of considering social and environmental context when studying aging in autism.

Although these findings should be considered preliminary, other psychosocial results supported this interpretation. Lower ratings of overall quality of life and health were associated with higher Brain SystemsAge, while greater loneliness was associated with higher Blood SystemsAge. These observations align with broader evidence that social connection, meaningful activity, and participation are important components of healthy aging, whereas loneliness, social isolation, chronic stress, and poor metabolic health are associated with less favorable later-life outcomes (53,54). Autistic adults experience disproportionately high rates of social isolation, unemployment or underemployment, and reduced community participation, making these factors particularly relevant to understanding aging in this population (30,55). Although these associations cannot establish directionality, they suggest that aging-related biological heterogeneity in autism may be shaped by potentially modifiable environmental factors.

Several limitations should be considered. The autistic sample was modest and restricted to a narrow age range, limiting power to detect age-dependent effects within autism and preventing direct inference about longitudinal aging trajectories. Controls were drawn from independent publicly available datasets, creating potential confounding between diagnostic group and data source despite joint preprocessing, matched tissue and array platform, technical sensitivity analyses, and leave-one-dataset-out analyses. Saliva is a heterogeneous peripheral tissue, and SystemsAge subscores should not be interpreted as direct measures of organ-specific aging. However, saliva offers practical advantages for inclusive autism research, and the strong association between measured BMI and DNAm BMI in the present sample provides evidence that biologically meaningful methylation-derived variation can be detected when cellular composition is appropriately accounted for (36,37). Finally, the within-autism analyses were exploratory; several measures were only available for a subset of participants, and some categorical comparisons involved small and uneven groups.

Longitudinal sampling across adulthood will determine whether these differences are stable, emerge progressively, or reflect genuinely accelerated trajectories, and whether they predict future health or functional decline. Larger cohorts combining epigenetic, genetic, and clinical measures may also help separate developmental liability from accumulated environmental influences, which are potentially modifiable.

In conclusion, altered aging-related biology in autism appeared system-specific rather than global, with the strongest differences concentrated in Brain and Blood SystemsAge. Within-autism variation was associated with metabolic and psychosocial factors rather than core clinical characteristics, suggesting that system-specific epigenetic measures may be useful for understanding heterogeneity and identifying pathways that support healthier aging in autistic adulthood.

## Supporting information

Supplementary Results

## Acknowledgments

We are extremely grateful to the participants and families who have taken part in the Early Diagnosis Study over three decades, and to everyone who has sustained the study across that period. We also thank the investigators who generated and publicly deposited the control methylation datasets analyzed here.

This work was supported by the National Institute of Mental Health (K01MH128612, to AD), the Eunice Kennedy Shriver National Institute of Child Health and Human Development (R01HD081199, to CL), and the National Institute of Mental Health (R01MH081873, to CL). The funders had no role in study design, data collection and analysis, decision to publish, or preparation of the manuscript.

## Data availability

Control DNA methylation data are publicly available through the NCBI Gene Expression Omnibus under accessions GSE151485, GSE232332, GSE111631, and GSE232891. Dataset selection, preprocessing, quality control, and calculation of all epigenetic aging measures are described in Methods S1 and S4 and Tables S1 to S3.

## References

1. Mason D, Stewart GR, Capp SJ, Happé F. Older Age Autism Research: A Rapidly Growing Field, but Still a Long Way to Go. Autism Adulthood. 2022;4(2):164–72. doi:10.1089/aut.2021.0041

2. Croen LA, Zerbo O, Qian Y, Massolo ML, Rich S, Sidney S, et al. The health status of adults on the autism spectrum. Autism. 2015;19(7):814–23. doi:10.1177/1362361315577517

3. Weir E, Allison C, Warrier V, Baron-Cohen S. Increased prevalence of non-communicable physical health conditions among autistic adults. Autism. 2021;25(3):681–94. doi:10.1177/1362361320953652

4. Hand BN, Angell AM, Harris L, Carpenter LA. Prevalence of physical and mental health conditions in Medicare-enrolled, autistic older adults. Autism. 2020;24(3):755–64. doi:10.1177/1362361319890793

5. Hirvikoski T, Mittendorfer-Rutz E, Boman M, Larsson H, Lichtenstein P, Bölte S. Premature mortality in autism spectrum disorder. Br J Psychiatry. 2016;208(3):232–8. doi:10.1192/bjp.bp.114.160192

6. O’Nions E, Lewer D, Petersen I, Brown J, Buckman JEJ, Charlton R, et al. Estimating life expectancy and years of life lost for autistic people in the UK: a matched cohort study. Lancet Reg Health – Eur. 2024;36:100776. doi:10.1016/j.lanepe.2023.100776

7. Harker SA, Al-Hassan L, Huentelman MJ, Braden BB, Lewis CR. APOE ε4-Allele in middle-aged and older autistic adults: associations with verbal learning and memory. Int J Mol Sci. 2023;24(21):15988.

8. Walsh MJM, Ofori E, Pagni BA, Chen K, Sullivan G, Braden BB. Preliminary findings of accelerated visual memory decline and baseline brain correlates in middle-age and older adults with autism: The case for hippocampal free-water. Front Aging Neurosci. 2022;14(1029166). doi:10.3389/fnagi.2022.1029166

9. Bessé M, Morel-Kohlmeyer S, Houy-Durand E, Prévost P, Tuller L, Bouazzaoui B, et al. Cognitive and Cerebral Aging Research in Autism: A Systematic Review on an Emerging Topic. Autism Res Off J Int Soc Autism Res. 2025 Jun;18(6):1122–45. doi:10.1002/aur.70031 PubMed PMID: 40196934; PubMed Central PMCID: PMC12166515.

10. Dickinson A, Ryan D, Carroll JE, Lord C. Aging in autism: A systematic review of cognitive, neural, and physical health findings. Neurosci Biobehav Rev. 2026 Sep;188:106820. doi:10.1016/j.neubiorev.2026.106820

11. Tse VW, Lei J, Crabtree J, Mandy W, Stott J. Characteristics of older autistic adults: A systematic review of literature. Rev J Autism Dev Disord. 2022;9(2):184–207.

12. Wang J, Christensen D, Coombes SA, Wang Z. Cognitive and brain morphological deviations in middle-to-old aged autistic adults: A systematic review and meta-analysis. Neurosci Biobehav Rev. 2024;163:105757. doi:10.1016/j.neubiorev.2024.105757

13. Bell CG, Lowe R, Adams PD, Baccarelli AA, Beck S, Bell JT, et al. DNA methylation aging clocks: challenges and recommendations. Genome Biol. 2019;20(1):249. doi:10.1186/s13059-019-1824-y

14. Horvath S, Raj K. DNA methylation-based biomarkers and the epigenetic clock theory of ageing. Nat Rev Genet. 2018;19:371–84. doi:10.1038/s41576-018-0004-3

15. Horvath S. DNA methylation age of human tissues and cell types. Genome Biol. 2013;14(10):R115. doi:10.1186/gb-2013-14-10-r115

16. Belsky DW, Caspi A, Corcoran DL, Sugden K, Poulton R, Arseneault L, et al. DunedinPACE, a DNA methylation biomarker of the pace of aging. eLife. 2022;11:e73420. doi:10.7554/eLife.73420

17. Levine ME, Lu AT, Quach A, Chen BH, Assimes TL, Bandinelli S, et al. An epigenetic biomarker of aging for lifespan and healthspan. Aging. 2018;10(4):573–91. doi:10.18632/aging.101414

18. Lu AT, Quach A, Wilson JG, Reiner AP, Aviv A, Raj K, et al. DNA methylation GrimAge strongly predicts lifespan and healthspan. Aging. 2019;11(2):303–27. doi:10.18632/aging.101684

19. Sehgal R, Markov Y, Qin C, Meer M, Hadley C, Shadyab AH, et al. Systems Age: a single blood methylation test to quantify aging heterogeneity across 11 physiological systems. Nat Aging. 2025;5(9):1880–96. doi:10.1038/s43587-025-00958-3

20. Harvanek ZM, Sehgal R, Borrus D, Kasamoto J, Priyanka A, Corley MJ, et al. Multidimensional Epigenetic Clocks Demonstrate Accelerated Aging Across Physiological Systems in Schizophrenia: A Meta-Analysis. medRxiv. 2024 Oct 30;2024.10.28.24316295. doi:10.1101/2024.10.28.24316295 PubMed PMID: 39574874; PubMed Central PMCID: PMC11581063.

21. Higgins-Chen AT, Boks MP, Vinkers CH, Kahn RS, Levine ME. Schizophrenia and Epigenetic Aging Biomarkers: Increased Mortality, Reduced Cancer Risk, and Unique Clozapine Effects. Biol Psychiatry. 2020;88(3):224–35. doi:10.1016/j.biopsych.2020.01.025

22. McKinney BC, Lin H, Ding Y, Lewis DA, Sweet RA. DNA methylation age is not accelerated in brain or blood of subjects with schizophrenia. Schizophr Res. 2018 Jun;196:39–44. doi:10.1016/j.schres.2017.09.025 PubMed PMID: 28988914; PubMed Central PMCID: PMC5886835.

23. Voisey J, Lawford BR, Morris CP, Wockner LF, Noble EP, Young RM, et al. Epigenetic analysis confirms no accelerated brain aging in schizophrenia. NPJ Schizophr. 2017 Sep 4;3:26. doi:10.1038/s41537-017-0026-4 PubMed PMID: 28871179; PubMed Central PMCID: PMC5583345.

24. Lord C, Risi S, DiLavore PS, Shulman C, Thurm A, Pickles A. Autism from 2 to 9 years of age. Arch Gen Psychiatry. 2006;63(6):694–701.

25. Lord C, McCauley JB, Pepa LA, Huerta M, Pickles A. Work, Living, and the Pursuit of Happiness: Vocational and Psychosocial Outcomes for Young Adults with Autism. Autism Int J Res Pract. 2020 Oct;24(7):1691–703. doi:10.1177/1362361320919246 PubMed PMID: 32431163; PubMed Central PMCID: PMC7541415.

26. Pickles A, McCauley JB, Pepa LA, Huerta M, Lord C. The adult outcome of children referred for autism: typology and prediction from childhood. J Child Psychol Psychiatry. 2020;61(7):760–7.

27. Lord C, Rutter M, DiLavore P, Risi S, Gotham K, Bishop S. Autism diagnostic observation schedule–2nd edition (ADOS-2). West Psychol Serv. 2012;284:474–8.

28. Rutter M, Le Couteur A, Lord C. Autism Diagnostic Interview-Revised (ADI-R). Los Angeles (CA): Western Psychological Services; 2003.

29. Elias R, Lord C. Diagnostic stability in individuals with autism spectrum disorder: insights from a longitudinal follow-up study. J Child Psychol Psychiatry. 2022;63(9):973–83. doi:10.1111/jcpp.13551

30. Anderson DK, Liang JW, Lord C. Predicting young adult outcome among more and less cognitively able individuals with autism spectrum disorders. J Child Psychol Psychiatry. 2014;55(5):485–94.

31. Asher SR, Hymel S, Renshaw PD. Loneliness in Children. Child Dev. 1984;55(4):1456–64. doi:10.2307/1130015

32. McConachie H, Mason D, Parr JR, Garland D, Wilson C, Rodgers J. Enhancing the Validity of a Quality of Life Measure for Autistic People. J Autism Dev Disord. 2018;48(5):1596–611. doi:10.1007/s10803-017-3402-z

33. The WHOQOL Group. Development of the World Health Organization WHOQOL-BREF Quality of Life Assessment. Psychol Med. 1998;28(3):551–8. doi:10.1017/S0033291798006667

34. Aryee MJ, Jaffe AE, Corrada-Bravo H, Ladd-Acosta C, Feinberg AP, Hansen KD, et al. Minfi: a flexible and comprehensive Bioconductor package for the analysis of Infinium DNA methylation microarrays. Bioinforma Oxf Engl. 2014 May 15;30(10):1363–9. doi:10.1093/bioinformatics/btu049 PubMed PMID: 24478339; PubMed Central PMCID: PMC4016708.

35. Min JL, Hemani G, Davey Smith G, Relton C, Suderman M. Meffil: efficient normalization and analysis of very large DNA methylation datasets. Bioinformatics. 2018;34(23):3983–9. doi:10.1093/bioinformatics/bty476

36. Raffington L, Tanksley PT, Sabhlok A, Vinnik L, Mallard T, King LS, et al. Socially Stratified Epigenetic Profiles Are Associated With Cognitive Functioning in Children and Adolescents. Psychol Sci. 2023 Feb 1;34(2):170–85. doi:10.1177/09567976221122760

37. Zarandooz S, Raffington L. Applying blood-derived epigenetic algorithms to saliva: cross-tissue similarity of DNA-methylation indices of aging, physiology, and cognition. Clin Epigenetics. 2025 Apr 23;17:61. doi:10.1186/s13148-025-01868-2 PubMed PMID: 40270051; PubMed Central PMCID: PMC12016411.

38. Zhou W, Laird PW, Shen H. Comprehensive characterization, annotation and innovative use of Infinium DNA methylation BeadChip probes. Nucleic Acids Res. 2017;45(4):e22. doi:10.1093/nar/gkw967

39. Fortin JP, Labbe A, Lemire M, Zanke BW, Hudson TJ, Fertig EJ, et al. Functional normalization of 450k methylation array data improves replication in large cancer studies. Genome Biol. 2014;15:503. doi:10.1186/s13059-014-0503-2

40. Middleton LYM, Dou J, Fisher J, Heiss JA, Nguyen VK, Just AC, et al. Saliva cell type DNA methylation reference panel for epidemiological studies in children. Epigenetics. 2022;17(2):161–77. doi:10.1080/15592294.2021.1890874

41. Horvath S, Oshima J, Martin GM, Lu AT, Quach A, Cohen H, et al. Epigenetic clock for skin and blood cells applied to Hutchinson Gilford Progeria Syndrome and ex vivo studies. Aging. 2018;10(7):1758–75. doi:10.18632/aging.101508

42. McCartney DL, Hillary RF, Stevenson AJ, Ritchie SJ, Walker RM, Zhang Q, et al. Epigenetic prediction of complex traits and death. Genome Biol. 2018 Sep 27;19:136. doi:10.1186/s13059-018-1514-1

43. Elliott HR, Tillin T, McArdle WL, Ho K, Duggirala A, Frayling TM, et al. Differences in smoking associated DNA methylation patterns in South Asians and Europeans. Clin Epigenetics. 2014 Feb 3;6(1):4. doi:10.1186/1868-7083-6-4

44. McCartney DL, Hillary RF, Conole ELS, Banos DT, Gadd DA, Walker RM, et al. Blood-based epigenome-wide analyses of cognitive abilities. Genome Biol. 2022;23:26. doi:10.1186/s13059-021-02596-5

45. Gentilini D, Cavagnola R, Possenti I, Calzari L, Ranucci F, Nola M, et al. Epigenetics of Autism Spectrum Disorders: A Multi-level Analysis Combining Epi-signature, Age Acceleration, Epigenetic Drift and Rare Epivariations Using Public Datasets. Curr Neuropharmacol. 2023;21(11):2362–73. doi:10.2174/1570159X21666230725142338 PubMed PMID: 37489793; PubMed Central PMCID: PMC10556384.

46. Liu L, Qi X, Cheng S, Meng P, Yang X, Pan C, et al. Epigenetic analysis suggests aberrant cerebellum brain aging in old-aged adults with autism spectrum disorder and schizophrenia. Mol Psychiatry. 2023 Nov;28(11):4867–76. doi:10.1038/s41380-023-02233-6

47. Okazaki S, Kimura R, Otsuka I, Funabiki Y, Murai T, Hishimoto A. Epigenetic clock analysis and increased plasminogen activator inhibitor-1 in high-functioning autism spectrum disorder. PLOS ONE. 2022 Feb 3;17(2):e0263478. doi:10.1371/journal.pone.0263478

48. Gilmore D, Hand BN. Diabetes mellitus in privately insured autistic adults in the United States. Autism. 2024;28(7):1785–94. doi:10.1177/13623613231206421

49. Starkstein S, Gellar S, Parlier M, Payne L, Piven J. High rates of parkinsonism in adults with autism. J Neurodev Disord. 2015;7(1):29. doi:10.1186/s11689-015-9125-6

50. Vivanti G, Tao S, Lyall K, Robins DL, Shea LL. The prevalence and incidence of early-onset dementia among adults with autism spectrum disorder. Autism Res. 2021;14(10):2189–99. doi:10.1002/aur.2590

51. Yin W, Reichenberg A, Schnaider Beeri M, Levine SZ, Ludvigsson JF, Figee M, et al. Risk of Parkinson Disease in Individuals With Autism Spectrum Disorder. JAMA Neurol. 2025 Jul 1;82(7):687–95. doi:10.1001/jamaneurol.2025.1284

52. Oblak L, van der Zaag J, Higgins-Chen AT, Levine ME, Boks MP. A systematic review of biological, social and environmental factors associated with epigenetic clock acceleration. Ageing Res Rev. 2021;69:101348. doi:10.1016/j.arr.2021.101348

53. Nakou A, Dragioti E, Bastas NS, Zagorianakou N, Kakaidi V, Tsartsalis D, et al. Loneliness, social isolation, and living alone: a comprehensive systematic review, meta-analysis, and meta-regression of mortality risks in older adults. Aging Clin Exp Res. 2025 Jan 21;37(1):29. doi:10.1007/s40520-024-02925-1

54. Polsky LR, Rentscher KE, Carroll JE. Stress-induced biological aging: A review and guide for research priorities. Brain Behav Immun. 2022;104:97–109. doi:10.1016/j.bbi.2022.05.016

55. Farley MA, McMahon WM, Fombonne E, Jenson WR, Miller J, Gardner M, et al. Twenty-year outcome for individuals with autism and average or near-average cognitive abilities. Autism Res. 2009;2(2):109–18. doi:10.1002/aur.69

