## Supplementary material for "System-Specific Epigenetic Aging Signatures in Autistic Adults": Table 1

*Sample Characteristics by Group in the Full Sample and Age- and Sex-Restricted Subsample*

|  | **Full Sample** | | |  | **Restricted Subsample** | | |
| --- | --- | --- | --- | --- | --- | --- | --- |
|  | **Autism**  **(N=37)** | **Controls**  **(N=188)** | ***p*** |  | **Autism**  **(N=31)** | **Controls**  **(N=32)** | ***p*** |
| **Age (years)** |  |  |  |  |  |  |  |
| n | 37 | 188 |  |  | 31 | 32 |  |
| Mean (SD) | 29.8 (1.1) | 36.8 (9.1) | < .001 |  | 29.8 (1.1) | 30.0 (3.6) | .697 |
| [Range] | [27.6–32.8] | [19–50] |  |  | [27.6–32.8] | [24–36] |  |
| **Estimated Cell-type Proportions** |  |  |  |  |  |  |  |
| n | 37 | 188 |  |  | 31 | 32 |  |
| Epithelial Cell Proportion, Mean (SD) | 0.05 (0.12) | 0.02 (0.09) | .182 |  | 0.06 (0.13) | 0.01 (0.03) | .122 |
| Immune Cell Proportion, Mean (SD) | 0.95 (0.12) | 0.98 (0.09) | .182 |  | 0.94 (0.13) | 0.99 (0.03) | .122 |
| **Sex** |  |  |  |  |  |  |  |
| n | 37 | 188 |  |  | 31 | 32 |  |
| Male, n (%) | 31 (83.8) | 97 (51.6) | < .001 |  | 31 (100) | 32 (100) | N/A |
| Female, n (%) | 6 (16.2) | 91 (48.4) |  |  | 0 | 0 |  |
| **Race/Ethnicity, n (%)** |  |  |  |  |  |  |  |
| n | 37 | 48 |  |  | 31 | 29 |  |
| Asian/Pacific Islander | 0 (0.0) | 3 (6.3) | 0.160 |  | 0 (0.0) | 3 (10.2) | 0.280 |
| Black or African American | 6 (16.2) | 10 (20.8) |  |  | 3 (9.7) | 5 (17.2) |  |
| Hispanic or Latino | 2 (5.4) | 4 (8.3) |  |  | 2 (6.5) | 0 |  |
| White or Caucasian | 29 (78.4) | 31 (64.6) |  |  | 26 (83.9) | 21 (72.4) |  |
| **Maternal education**  **≥4-year degree, n (%)** |  |  |  |  |  |  |  |
| n | 36 | — | N/A |  | 31 | — | N/A |
| Yes | 24 (66.7) |  |  |  | 20 (64.5) |  |  |
| No | 12 (33.3) |  |  |  | 11 (35.5) |  |  |

***Note.*** Age refers to age at saliva collection. The age- and sex-restricted subsample included autistic males and male controls aged 24–36 years. Race/ethnicity data were available for 48 controls in the full sample and 29 controls in the restricted subsample. Maternal education was unavailable for one autistic participant. Group differences were assessed using Fisher’s exact tests or χ² tests for categorical variables, depending on expected cell counts, and independent-samples t tests or Wilcoxon rank-sum tests for continuous variables, as appropriate.
