## Supplementary material for "System-Specific Epigenetic Aging Signatures in Autistic Adults": Table 2

*Clinical Characteristics of the Autistic Sample*

|  | **n** | **M (SD) or n (%)** | **[Range]** |
| --- | --- | --- | --- |
| **Age of saliva collection (years)** | 37 | 29.8 (1.1) | [27.6–32.8] |
| **Age at initial autism diagnosis (years)** | 37 | 5.23 (5.53) | [1.33–25.8] |
| **IQ grouping, n (%)** |  |  |  |
| VIQ ≥ 70 |  | 22 (59.5) |  |
| VIQ < 70 |  | 15 (40.5) |  |
| **Cognitive assessment (closest to saliva collection)** |  |  |  |
| Age at Assessment (Years) | 37 | 21.2 (4.2) | [8.2–29.2] |
| Full Scale IQ (FSIQ) | 37 | 73.6 (44.6) | [6–130] |
| Verbal IQ (VIQ) | 37 | 72.8 (46.9) | [4–139] |
| Non-verbal IQ (NVIQ) | 37 | 72.8 (42.4) | [5–133] |
| **ADOS-2 (closest to saliva collection)** |  |  |  |
| Age at assessment (years) | 37 | 23.7 (5.6) | [8.1–30.1] |
| CSS Social Affect | 37 | 6.0 (2.0) | [2–9] |
| CSS RRB | 37 | 6.1 (2.3) | [1–10] |
| **BMI (closest to saliva collection)** |  |  |  |
| Age at assessment (years) | 34 | 20.5 (3.27) | [12.7–28.2] |
| BMI (kg/m^2^) | 34 | 28.3 (9.0) | [16.3–51.7] |
| BMI < 25, n (%) |  | 15 (44.1) |  |
| BMI ≥25, n (%) |  | 19 (55.9) |  |
| **Asher Loneliness** |  |  |  |
| Age at assessment (years) | 16 | 27.7 (3.7) | [21.7–31.4] |
| Asher Total Score | 16 | 38.9 (10.5) | [18-53] |
| **QOL (WHOQOL-BREF)** |  |  |  |
| All respondents | 28 | 4.07 (0.72) | [2-5] |
| Self-report | 11 | 3.91 (1.04) | [2-5] |
| Caregiver report | 24 | 4.17 (0.48) | [3-5] |

***Note.*** Values are M (SD) with observed ranges unless otherwise indicated. Cognitive, ADOS-2, BMI, WHOQOL-BREF, and Asher assessments were selected from the assessment closest to saliva collection for each participant. BMI data were unavailable for three participants. The proportion in the normal-weight-or-below range was higher in this subsample than in the broader EDX cohort (44.1% vs. 31%; Byrne et al., 2022). VIQ groups were defined as VIQ ≥ 70 and VIQ < 70. The Asher Loneliness and Social Dissatisfaction Scale was administered by self-report only; all 16 participants with Asher data had VIQ ≥ 70. Of the 28 participants with WHOQOL-BREF data, 11 had self-report data and 24 had caregiver-report data; seven had both. When both were available, the assessment closest to saliva collection was used for the ‘all respondents’ score. BMI = body mass index; FSIQ = full-scale IQ; VIQ = verbal IQ; NVIQ = nonverbal IQ; ADOS-2 = Autism Diagnostic Observation Schedule, Second Edition; CSS = calibrated severity score; SA = Social Affect; RRB = Restricted and Repetitive Behavior; WHOQOL-BREF = World Health Organization Quality of Life–Brief; QOL = quality of life.
