## Supplementary material for "System-Specific Epigenetic Aging Signatures in Autistic Adults": Table 3

**Table 3.** Group Differences in Epigenetic Aging Measures in the Full Sample and Age- and Sex-Restricted Subsample

|  | **Full Sample** | | | |  | **Restricted Subsample** | | | |
| --- | --- | --- | --- | --- | --- | --- | --- | --- | --- |
| Clock | β (std) | 95% CI | *P* | *FDR q* |  | β (std) | 95% CI | *P* | *FDR q* |
| Horvath Skin & Blood | 0.01 | [-0.11, 0.14] | .824 | .875 |  | -0.09 | [-0.32, 0.15] | .467 | .582 |
| GrimAge | 0.03 | [-0.15, 0.20] | .781 | .875 |  | 0.17 | [-0.14, 0.48] | .288 | .480 |
| Dunedin PACE | 0.02 | [-0.33, 0.40] | .875 | .875 |  | 0.33 | [-0.28, 0.93] | .284 | .480 |
| Systems Age | 0.21 | [-0.11, 0.53] | .202 | .346 |  | 0.27 | [-0.15, 0.69] | .207 | .480 |
| Blood | **0.65** | **[0.31, 0.99]** | **< .001** | **.002** |  | **0.63** | **[0.24, 1.01]** | **.002** | **.021** |
| Brain | **0.81** | **[0.44, 1.19]** | **< .001** | **< .001** |  | **0.61** | **[0.22, 1.01]** | **.003** | **.021** |
| Heart | 0.27 | [-0.07, 0.61] | .120 | .258 |  | 0.25 | [-0.16, 0.66] | .233 | .480 |
| Hormone | -0.04 | [-0.34, 0.26] | .791 | .875 |  | 0.14 | [-0.31, 0.59] | .544 | .582 |
| Immune | 0.03 | [-0.26, 0.32] | .852 | .875 |  | 0.17 | [-0.26, 0.60] | .434 | .582 |
| Inflammation | 0.20 | [-0.11, 0.52] | .208 | .346 |  | 0.16 | [-0.28, 0.61] | .469 | .582 |
| Kidney | -0.25 | [-0.53, 0.04] | .088 | .258 |  | -0.26 | [-0.70, 0.18] | .236 | .480 |
| Liver | **0.58** | **[0.22, 0.94]** | **.002** | **.008** |  | 0.32 | [-0.08, 0.71] | .117 | .439 |
| Lung | -0.06 | [-0.36, 0.24] | .698 | .875 |  | -0.01 | [-0.44, 0.43] | .974 | .974 |
| Metabolic | 0.38 | [0.03, 0.73] | .035 | .130 |  | 0.35 | [-0.07, 0.77] | .100 | .439 |
| Musculoskeletal | 0.27 | [-0.05, 0.59] | .104 | .258 |  | 0.17 | [-0.36, 0.71] | .523 | .582 |

**Note.** Standardized β estimates represent the adjusted difference between autistic and non-autistic participants in control-group standard deviation units, with positive values indicating higher scores in the autism group. Full-sample models adjusted for chronological age, sex, and estimated epithelial cell proportion. The age- and sex-restricted subsample included autistic males and male controls aged 24–36 years. FDR q values were calculated across all 15 epigenetic measures within each analytic sample using the Benjamini–Hochberg procedure. Bold values indicate findings that remained significant after FDR correction (q < .05).
