## Supplemental Table 1 for "System-Specific Epigenetic Aging Signatures in Autistic Adults"

**Supplementary Table S1.** GEO Control Datasets: Characteristics and Quality Control Summary

| **Dataset Label** | **GEO Dataset** | **Study Purpose** | **Controls Included** | **Tissue;**  **Platform** | **Original sample** | | | **QC Exclusions** | **Final sample** | | | **Cell Composition Immune Cells Mean (SD)** |
| --- | --- | --- | --- | --- | --- | --- | --- | --- | --- | --- | --- | --- |
|  |  |  |  |  | **N** | **Age, Mean (SD) [Range]** | **M/F** |  | **N** | **Age, Mean (SD) [Range]** | **M/F** |  |
| Control Dataset 1 | GSE151485 | Examined effects of short-term opioid exposure on DNAm. | Pre-opioid baseline samples used as controls. | Saliva;  EPIC v1 | 31 | 28.3 (7.59) [19–48] | 14/17 | 0 | 31 | 28.3 (7.59) [19–48] | 14/17 | 0.936 (0.202) |
| Control Dataset 2 | GSE232332 | Investigated DNAm changes associated with esophageal cancer relative to healthy controls. | Healthy control samples only. | Saliva;  EPIC v1 | 38 | 40.0 (7.73) [24–50] | 26/12 | 4 [3 excluded for detection p-value threshold; 1 excluded for sex mismatch] | 34 | 39.7 (7.85) [24–50] | 22/12 | 0.994 (0.023) |
| Control Dataset 3 | GSE111631 | Genome-wide DNAm profiling of normal saliva from males and females. | All samples annotated as healthy controls included. | Saliva;  EPIC v1 | 18 | 43.1 (6.36) [29–50] | 16/2 | 1 [1 excluded for detection p-value threshold] | 17 | 43.4 (6.36) [29–50] | 15/2 | 1.00 (0.00) |
| Control Dataset 4 | GSE232891 | Epigenetic profiling of saliva in Crohn’s disease and ulcerative colitis with healthy controls. | Healthy controls (no Crohn’s disease or ulcerative colitis) included. | Saliva;  EPIC v1 | 117 | 36.9 (8.68) [21–50] | 54/63 | 11 [2 excluded for detection p-value threshold; 1 excluded for bead count; 8 excluded for sex mismatch] | 106 | 37.3 (8.57) [21–50] | 46/60 | 0.985 (0.046) |
|  |  |  |  | **Total:** | **204** | **37.5 (8.54) [19–50]** | **110/94** | 16 | 188 | **36.8 (9.1) [19–50]** | **97/91** | **0.983 (0.061)** |

Note. M/F = male/female; EPIC = Illumina HumanMethylationEPIC v1.0 BeadChip. All datasets comprised saliva samples assayed using the EPIC v1 array. Cell composition was estimated from DNA methylation data using EpiDISH and is reported as the estimated immune cell proportion. QC exclusions were based on detection p values, bead count, and disagreement between methylation-predicted and reported sex, as applicable
