## Supplemental Table 2 for "System-Specific Epigenetic Aging Signatures in Autistic Adults"

**Supplementary Table S2.** Probe-Level Quality Control: Exclusion Criteria and Removal Counts

| **Category** | **Description** | **Threshold** | **Probes Removed** | **% of EPIC Array** |
| --- | --- | --- | --- | --- |
| Detection p-value | Probes with detection p-value > 0.01 in > 5% of samples | p > 0.01 | 4189 | 0.48 |
| Bead count | Probes with < 3 beads in > 1% of samples | < 3 beads | 2900 | 0.33 |
| **Quality Probes (Both Categories above)** |  |  | 6905 | 0.80 |
| Sex chromosomes | Probes mapping to chromosomes X or Y | — | 19627 | 2.27 |
| SNP overlap | Probes with CpG site or single-base extension overlapping a known SNP | — | 30435 | 3.52 |
| **Sex and SNP** |  |  | 49733 | 5.74 |
| **Quality Probes (det p and bead count) + Sex and SNP** |  |  | 54906 | 6.34 |
| Cross-reactive (sensitivity analysis only) | Probes on the Zhou et al. (2017) cross-reactive probe list; excluded in sensitivity analyses only | — | 65537 | 7.57 |
| **Quality Probes (det p and bead count) + Sex and SNP+ Cross reactive** |  |  | 114306 | 13.20 |
| **Total probes removed** | **74,122** | **6.9%** |  |  |

Note. Counts reflect probes removed from the full EPIC array (865,859 probes present across all samples after import). Categories are not mutually exclusive; the total reflects the union of all excluded probes. Cross-reactive probes were excluded only in sensitivity analyses and are not included in the total. Primary analyses were conducted on the remaining 791,737 probes after applying the first four exclusion criteria.
