## Supplemental Table 3 for "System-Specific Epigenetic Aging Signatures in Autistic Adults"

**Supplementary Table S3.** Epigenetic Measures Implementation Details and CpG Probe Coverage

| **Measure** | **Category** | **R Implementation** | **Reference** | **CpGs Required** | **Primary Analysis** | | **Sensitivity Analysis (excl. cross-reactive probes)** | |
| --- | --- | --- | --- | --- | --- | --- | --- | --- |
|  |  |  |  |  | **N CpGs Present** | **Coverage (%)** | **N CpGs Present** | **Coverage (%)** |
| **Epigenetic Clocks** | | | | | | | | |
| Horvath SkinBlood | First-generation clock | methylclock (R package) | Horvath (2018) Aging | 391 | 391 | 100.0 | 385 | 98.5 |
| GrimAge | Second-generation clock | meffonym (R package) | Lu et al. (2019) Nature Aging | 1030 | 1022 | 99.2 | 865 | 83.9 |
| DunedinPACE | Third-generation clock | DunedinPACE (R package) | Belsky et al. (2022) eLife | 173 | 173 | 100.0 | 172 | 99.4 |
| **Systems Age Scores** | | | | | | | | |
| Systems Age (Composite & Organ Specific) | Organ-specific aging | methylCIPHER (R package) | Sehgal et al. (2025) Nature Aging | 125175 | 124605 | 99.5 | 119267 | 95.3 |
| **Epigenetic Covariates** | | | | | | | | |
| DNAm BMI | Lifestyle covariate | methylCIPHER (R package); McCartney et al. weights | McCartney et al. (2018) Int J Epidemiology | 1109 | 1058 | 95.40 | 1058 | 95.40 |
| DNAm Smoking | Lifestyle covariate | EpiSmokEr (R package); Elliott et al. CpGs | Elliott et al. (2014) Clin Epigenetics | 187 | 172 | 91.97 | 172 | 91.97 |
| DNAm Alcohol | Lifestyle covariate | methylCIPHER (R package); McCartney et al. weights | McCartney et al. (2018) Int J Epidemiology | 450 | 427 | 94.88 | 427 | 94.88 |

*Note. Primary analysis used functionally-normalized β-values with removal of probes failing detection p-value or bead count thresholds, sex-chromosome probes, and SNP-overlapping probes. Sensitivity analysis additionally excluded cross-reactive probes (Zhou et al., 2017). Coverage = percentage of algorithm-required CpG sites present in the dataset after QC filtering. Systems Age sub-scores (Brain, Lung, Musculoskeletal, etc.) share the same CpG set as the composite Systems Age score; system-specific scores are derived from weighted principal components rather than direct CpG contributions. [xx] = Abby add.*
