## Supplementary Figure 1 for "System-Specific Epigenetic Aging Signatures in Autistic Adults"

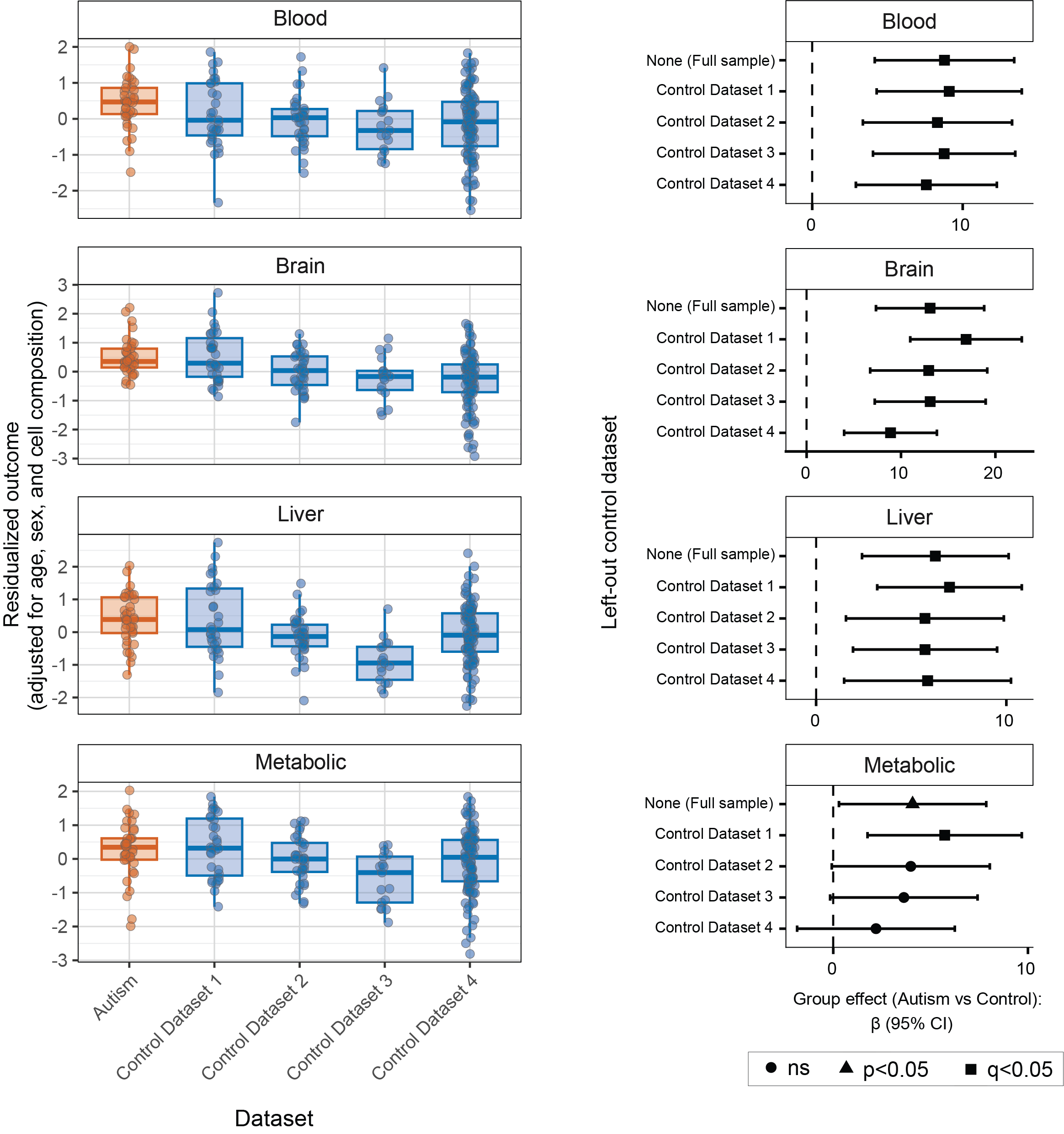

**Supplementary Figure 1.** Leave-one-dataset-out (LODO) analyses for Brain, Blood, Liver, and Metabolic subscores. Left: Residualized scores for the autistic group (red) and each individual control dataset (blue). Right: Modelled group differences between autistic and control participants (β, 95% CI) after sequentially excluding each control dataset. The axis labels indicate which control dataset was excluded from the model; “None” represents the primary model reported in the main analyses, in which all control datasets were included.
