## Supplementary Methods for "System-Specific Epigenetic Aging Signatures in Autistic Adults"

S1. Obtaining Control Data

A systematic search of the NCBI Gene Expression Omnibus (GEO) was conducted on October 27, 2025, to identify publicly available DNA methylation datasets. The search query ((saliva) AND idat) AND “Homo sapiens” was used, with study type filters applied to restrict results to methylation profiling by SNP array, array, genome tiling array, or high-throughput sequencing. This search yielded 42 datasets.

Each dataset was screened according to five pre-specified inclusion criteria: (1) availability of raw IDAT files for download; (2) methylation profiling performed using the Illumina HumanMethylationEPIC (EPIC v1.0) BeadChip; (3) saliva as the reported biological tissue; (4) inclusion of adult participants (aged 18 years or older); and (5) metadata containing at least participant age and sex. Datasets failing any criterion were excluded. Common exclusion reasons included pediatric samples, use of non-EPIC array platforms, missing required metadata, or analysis of tissues other than saliva.

Four datasets (GSE151485, GSE232332, GSE111631, GSE232891) satisfied all inclusion criteria and were incorporated into the control sample pool. Within each dataset, only participants classified as healthy or typically developing by the original investigators were included. Individuals with neurological, psychiatric, or major medical conditions were excluded. Participants aged 50 or older were also excluded to better align with the autistic cohort's age range.

Dataset-level details, including sample sizes before and after quality control, are presented in Supplementary Table S1. Note. Control ages were available only to the nearest year; therefore, ages were rounded to the nearest year for both groups in analyses involving controls, although autistic participants’ ages are presented at their original precision in the table.

S2. Co-occurring Conditions

Initial autism diagnoses were made at the age-2 wave for n=25 participants (67.6%), the age-3 wave for n=1 (2.7%), the age-5 wave for n=3 (8.1%), the age-9 wave for n=5 (13.5%), and the age-18 wave for n=3 (8.1%). The median age at initial diagnosis was 2.83 years (mean = 5.23 years).

To characterize adult co-occurring conditions, diagnoses recorded from Wave 25 onwards (minimum participant age = 22.8 years) were considered. Co-occurring diagnostic information was available for 36 of 37 participants. Participants contributed a median of 6 distinct assessment timepoints reporting diagnostic and medication data (mean = 5.94; range = 1–13). The average age across participants’ assessment timepoints was 28.66 years. Participants were, on average, 26.38 years old at their first included assessment and 30.96 years old at their last. The median follow-up period was 5.45 years (mean = 4.58; range = 0–6.31 years). Participants were classified as having a given condition if it was reported at any visit during this period.

Twenty-eight participants (77.8% of those with available data) had at least one co-occurring diagnosis. Co-occurring conditions frequently overlapped; diagnostic prevalences and overlap are presented in Table S7. Examples of specific diagnoses included in each category are included below.

Mood/anxiety disorders: anxiety, depression, obsessive-compulsive disorder (OCD), panic disorder, bipolar disorder, mood disorder, and catatonia.

Behavioral disorders: aggression.

ADHD/ODD: attention-deficit/hyperactivity disorder (ADHD), oppositional defiant disorder (ODD), and hyperactivity.

Epilepsy/seizure disorders: epilepsy and seizure disorders.

Other physical health conditions: type I/II diabetes, hypertension, thyroid disorders, hyperlipidemia, brain cyst, lupus, and kidney stones.

Note. Other physical health conditions comprised a heterogeneous range of nonbehavioral and nonpsychiatric conditions, distinct from behavioral disorder, epilepsy/seizures, ODD/ADHD, and mood/anxiety disorder. Thirteen participants had at least one condition in this category, including monocular blindness (n = 1), asthma (n = 1), hormonal or thyroid conditions (n = 2), kidney problems or kidney stones (n = 1), hearing loss (n = 1), hyperlipidemia or high cholesterol (n = 3), hypertension (n = 3), a brain cyst (n = 1), prediabetes (n = 1), lupus (n = 1), type 1 diabetes (n = 1), and type 2 diabetes (n = 1). Counts refer to conditions and are not mutually exclusive, as some participants had more than one physical health condition.

S3. Medication Use

Medication histories recorded from Wave 25 onwards (minimum participant age = 22.8 years) were reviewed. Medication data were available for 36 of 37 participants. Participants were classified as having a history of neuroactive medication use if they reported taking a centrally acting psychotropic or neuroactive medication at any assessment during this period.

Of the 36 participants with available data, 26 (72.2%) had a history of neuroactive medication use. Medication status was coded as a binary variable; specific medications included in each category are detailed below.

Antipsychotics: aripiprazole, olanzapine, quetiapine, chlorpromazine.

Antidepressants: fluoxetine, sertraline, escitalopram, citalopram, paroxetine, venlafaxine, mirtazapine, trazodone.

Anxiolytic/sedative medications: lorazepam, clonazepam, buspirone, hydroxyzine.

Anticonvulsants: levetiracetam, oxcarbazepine, lamotrigine, valproate, cannabidiol.

Stimulant/ADHD medications: amphetamine salts and methylphenidate.

Non-neuroactive medications and dietary supplements were excluded from the binary medication variable. Examples included levothyroxine, insulin and other antidiabetic medications, rosuvastatin, bisoprolol, antihistamines, respiratory medications, melatonin, vitamins, acetyl-L-carnitine, and coenzyme Q10.

S4. Preprocessing and Quality Control

Preprocessing and quality control (QC) were conducted using the minfi (v1.46.0)(Aryee et al., 2014), meffil (v1.3.6) (Min et al., 2018), EpiDISH (v2.16.0) (Teschendorff et al., 2017), and tidyverse (v2.0.0) (Wickham et al., 2019) packages in R (version 4.5.1). IDAT files were imported using minfi::read.metharray.exp(). Probes present across all samples were initially retained, yielding 865,858 CpG sites for downstream filtering.

Sample-Level QC. Samples were excluded if they met any of the following criteria: (i) >1% of probes with detection p-value >0.01 (minfi::detectionP), (ii) log-transformed median methylated and unmethylated signal intensity <8.5 (minfi::getQC), (iii) >1% of probes with bead count <3 (meffil), or (iv) predicted sex derived from X-chromosome methylation patterns (minfi::getSex) did not match reported sex. Sixteen samples (all from control datasets) were excluded following these criteria; per-criterion exclusion counts are reported in Supplementary Table S1. After quality control, the final analytic sample comprised 37 autistic and 188 control participants (Table 1).

Probe-Level QC. Probes were excluded if they: (i) had detection p-values > 0.01 in more than 1% of samples; (ii) had bead counts < 3 in more than 1% of samples; (iii) mapped to sex chromosomes; or (iv) overlapped known single-nucleotide polymorphisms (SNPs). Probe-level filtering was applied after normalization to ensure all probes contributed to background correction prior to exclusion. Together, these criteria resulted in the removal of 54,906 probes (6.34%), leaving 810,952 probes for primary analyses. Cross-reactive probes (Zhou et al., 2017) were additionally excluded in sensitivity analyses (Section 2.5.3). Detailed probe removal counts by criterion are provided in Supplementary Table S2.

Normalization and cell-type estimation. Methylation data were normalized using functional normalization (minfi::preprocessFunnorm), which uses within-array control probe intensities to remove technical variation prior to quantile normalization and does not require knowledge of sample group labels (Fortin et al., 2014). This approach is well-suited to multi-cohort data, in which case-control status is correlated with data source, as it reduces between-array and between-batch technical variation while preserving biological signal. Cell-type composition was estimated using the EpiDISH package with the EPIC-based bead-sorted saliva reference panel (EH4540) (Middleton et al., 2022). Estimated cell-type proportions were included as covariates in all regression models to account for cellular heterogeneity inherent to saliva-derived samples.

S5. Analyses.

Categorical predictors were analyzed as adjusted group differences and were restricted to comparisons with at least five participants per group. Where necessary because of sample-size constraints, variables were collapsed into binary categories. Race and ethnicity were coded as non-Hispanic White versus all other racial and ethnic groups. Maternal education was coded as completion of at least four years of college versus fewer than four years.

Continuous predictors included IQ, ADOS-2 calibrated severity scores, BMI, loneliness, and quality of life. When predictor assessments were obtained at a different time from saliva collection, models additionally adjusted for the interval between assessment and saliva collection. Given the modest sample size and exploratory nature of these analyses, findings were interpreted with emphasis on effect sizes, confidence intervals, and consistency across models.
