## Supplementary Results for "System-Specific Epigenetic Aging Signatures in Autistic Adults"

S1. Additional age-association analyses

Within the autistic cohort, the estimated slopes remained positive and were similar in magnitude, but the associations were less precise. Chronological age was significantly associated with Horvath skin-and-blood age (r = .38, b = 1.01, SE = 0.43, 95% CI [0.14, 1.88], p = .025). Associations with GrimAge (r = .31, b = 0.76, SE = 0.40, 95% CI [−0.06, 1.58], p = .067) and the SystemsAge composite (r = .22, b = 1.68, SE = 1.32, 95% CI [−1.00, 4.37], p = .211) were in the same direction but did not reach statistical significance. The wider confidence intervals and lower correlations likely reflect the smaller sample size and substantially narrower age range within the autistic subgroup.

S2. Sensitivity Analyses

In sensitivity analyses using noob background-corrected data and additionally adjusting for the first 10 technical principal components, the Brain and Blood SystemsAge group differences remained significant after FDR correction (Brain: p < .001, q < .001; Blood: p < .001, q = .005; Supplementary Table S4). The Liver score difference was not replicated (p = .360), and the Metabolic score difference remained nonsignificant (p = .070).

The Brain and Blood SystemsAge group differences were robust across all alternative analytic specifications (Figure 2C). Both remained significant after FDR correction in the age- and sex-restricted subsample (Brain: β = 0.74, 95% CI [0.26, 1.21], q = .013; Blood: β = 0.59, 95% CI [0.22, 0.95], q = .012), after exclusion of cross-reactive probes (Brain: β = 0.82, 95% CI [0.44, 1.19], q < .001; Blood: β = 0.63, 95% CI [0.30, 0.97], q = .001), and after additional adjustment for DNAm BMI, DNAm smoking, or both together (Brain: β = 0.77–0.82, all q < .001; Blood: β = 0.61–0.68, all q ≤ .002). In the most extensively adjusted model, which additionally included a DNAm measure of general cognitive ability (DNAm-g), both effects were attenuated but remained significant (Brain: β = 0.45, 95% CI [0.13, 0.76], q = .045; Blood: β = 0.42, 95% CI [0.11, 0.72], q = .045).

The Liver SystemsAge difference was similarly robust across analytic variations (β = 0.54–0.63, all q ≤ .007) but was not significant in the age- and sex-restricted subsample (β = 0.34, 95% CI [−0.09, 0.76], q = .351), and fell short of significance in the most extensively adjusted model (β = 0.42, 95% CI [0.09, 0.75], q = .052). The Metabolic difference was nonsignificant in the primary model but reached significance after adjustment for DNAm smoking (β = 0.42, 95% CI [0.07, 0.77], q = .035) and for DNAm smoking and BMI together (β = 0.38, 95% CI [0.06, 0.70], q = .040).

Composite aging measures (Horvath, GrimAge, DunedinPACE) remained nonsignificant across sensitivity analyses, supporting a pattern of group differences concentrated in specific physiological systems rather than generalized across epigenetic aging measures.

*Control sample composition****:*** Leave-one-dataset-out analyses indicated that the Brain, Blood, and Liver group differences were not driven by any single control dataset. For all three measures, coefficients remained positive, confidence intervals excluded zero, and differences remained significant after FDR correction in every iteration (Brain: *b* = 8.89–16.9, all *q* < .001; Blood: *b* = 7.58–9.10, all *q* ≤ .002; Liver: *b* = 5.72–7.01, all *q* ≤ .009; Supplementary Figure S1). Estimates were smallest when Control Dataset 4 was excluded, the iteration with the smallest remaining control sample (*n* = 82).

The Metabolic difference was less consistent. It was nominally significant in the full sample but did not survive FDR correction (*b* = 4.08, 95% CI [0.30, 7.85], *p* = .035, *q* = .075), and reached significance only when Control Dataset 1 was excluded (*b* = 5.72, 95% CI [1.76, 9.68], *q* = .024). Exclusion of the remaining control datasets yielded nonsignificant estimates (*q* = .075–.285), indicating that Metabolic SystemsAge differences were sensitive to control sample composition.

S3. Structured Daytime Engagement Follow-up Analyses

To examine whether associations with structured daytime engagement were specific to employment or education, occupied participants were further separated into those engaged in work or education and those engaged in other structured daytime activities.

For Blood SystemsAge, there was an overall difference across the three groups (omnibus p = .034). Participants who were not occupied had higher Blood SystemsAge than both those engaged in work or education (β = 0.74, 95% CI [0.11, 1.38], p = .023) and those engaged in other structured activities (β = 0.98, 95% CI [0.23, 1.74], p = .012). The work/education and other-activity groups did not differ significantly (β = 0.23, 95% CI [−0.30, 0.76], p = .383).

A similar pattern was observed for Brain SystemsAge (omnibus p = .005). Participants who were not occupied had higher Brain SystemsAge than both the work/education group (β = 1.92, 95% CI [0.76, 3.09], p = .002) and the other-activity group (β = 1.55, 95% CI [0.34, 2.76], p = .014). There was little evidence of a difference between the two occupied groups (β = −0.42, 95% CI [−0.92, 0.07], p = .091).

Metabolic SystemsAge showed the same pattern (omnibus p = .001). Participants who were not occupied had higher Metabolic SystemsAge than both the work/education group (β = 0.95, 95% CI [0.37, 1.54], p = .002) and the other-activity group (β = 1.39, 95% CI [0.67, 2.11], p < .001), while the two occupied groups did not differ significantly (β = 0.42, 95% CI [−0.16, 1.01], p = .151).

Evidence was weaker for Liver SystemsAge (omnibus p = .079). Liver SystemsAge was higher among participants who were not occupied than among those engaged in other structured activities (β = 1.16, 95% CI [0.11, 2.21], p = .031), but not significantly higher than among those engaged in work or education (β = 0.80, 95% CI [−0.21, 1.82], p = .117). The two occupied groups did not differ significantly (β = 0.32, 95% CI [−0.24, 0.88], p = .247).

Overall, these exploratory analyses were consistent with the primary two-group findings and suggested that the associations with Brain and Metabolic SystemsAge, and less consistently Blood SystemsAge, were related to structured daytime engagement more broadly rather than to employment or education specifically.

S4. Quality-of-Life Sensitivity Analyses

Quality of life was assessed using WHOQOL-BREF ratings of overall quality of life and overall health (n = 28; Supplementary Table S10). Lower ratings on both measures were associated with higher Brain SystemsAge (overall quality of life: β = −0.44, 95% CI [−0.81, −0.06], p = .024; overall health: β = −0.43, 95% CI [−0.70, −0.17], p = .003). Lower overall quality of life was also associated with higher Metabolic SystemsAge (β = −0.48, 95% CI [−0.87, −0.09], p = .019).

When analyses were restricted to caregiver-report data (n = 24), the associations with Brain SystemsAge remained significant for both overall quality of life (β = −0.37, 95% CI [−0.73, −0.01], p = .047) and overall health (β = −0.44, 95% CI [−0.79, −0.09], p = .016). The Metabolic SystemsAge association was weaker and did not reach significance in the caregiver-report analysis (β = −0.41, 95% CI [−0.83, 0.02], p = .061).

In fully adjusted models, the associations between Brain SystemsAge and both overall quality of life (β = −0.41, 95% CI [−0.70, −0.12], p = .008) and overall health (β = −0.40, 95% CI [−0.63, −0.16], p = .002) remained significant. In contrast, the Metabolic SystemsAge association with overall quality of life was attenuated and no longer significant (β = 0.06, 95% CI [−0.38, 0.51], p = .765).
